# Presaccadic modulation of lateral interactions

**DOI:** 10.1101/2025.02.10.637412

**Authors:** Gabriela Mueller de Melo, Isabella de Oliveira Pitorri, Gustavo Rohenkohl

## Abstract

Lateral interactions are pervasive in early visual processing, contributing directly to processes such as object grouping and segregation. This study examines whether saccade preparation - known to affect visual perception - modulates lateral interactions. In a psychophysical task, participants were instructed to detect a Gabor target flanked by two adjacent Gabors, while they either prepared a saccade to the target or maintained central fixation. Flanker gratings could be iso- or orthogonally oriented to the target and were positioned at three different distances (4λ, *8*λ, and 16λ). Contrast thresholds for target detection were estimated in each condition using a 3-down/1-up staircase procedure. The results showed that in both presaccadic and fixation conditions, the target was suppressed at the shortest flanker distance (4λ), revealed by markedly higher thresholds in iso-oriented compared to orthogonal flanker configurations. Lateral interaction effects were completely abolished at their largest separation (16λ. Interestingly, at the intermediate flanker distance (8λ), we observed an increase in suppression of targets presented during the presaccadic period, but not in the fixation condition. This result suggests that saccade preparation can modulate lateral interactions, promoting suppressive effects over larger distances. These findings are consistent with the visual remapping phenomenon observed before saccade execution, especially the convergent remapping of receptive fields in oculomotor and visual areas. Finally, this presaccadic expansion of inhibitory lateral interactions could assist target selection by suppressing homogeneous peripheral signals - such as iso-oriented collinear patterns - while prioritizing the processing of more salient visual information.

## INTRODUCTION

Contextual modulations in vision refer to the enhancement or suppression of a stimulus based on interactions with other stimuli in its surroundings^[1]^. In the primary visual cortex (V1), the response of a neuron to stimulation within the classical receptive field (CRF) is highly sensitive to signals coming from the surrounding area^[2]^. Thus, rather than operating exclusively as local feature detectors within the CRF, V1 neurons can integrate visual information conveyed from the CRF surrounds^[3]^. A common example of contextual modulation is the effect of lateral interactions, observed as changes in visual sensitivity to central targets when adjacent flanking elements are introduced^[4]^.

For lateral interactions in the center of the visual field, psychophysical studies have shown that target sensitivity is suppressed by iso-oriented collinear flankers in close proximity, and facilitated when the flankers are placed further away from the target^[4–6]^. In peripheral vision, the effect of facilitation by iso-oriented flankers is less robust^[6–8]^, with significant differences from central vision already present at eccentricities of 1-2 degrees of visual angle (dva)^[9]^. In turn, iso-orientation suppression prevails over a wider range of target-to-flanker distances in the near-periphery, around 3-4 dva of eccentricity^[7;8;10;11]^. Similar suppressive^[12]^ and facilitatory^[13]^ effects have been described for neuronal responses in area V1.

Multiple neural mechanisms - including horizontal, feedback, and feedforward connections - are known to mediate contextual modulations within V1^[14]^. Although spatial interactions between nearby stimuli can be explained by horizontal connections, interactions over longer distances are thought to depend on feedback from extrastriate areas, which convey signals from a larger area of visual space^[3]^. Feedback control is thought to operate in V1 through highly flexible mechanisms^[15]^, adjusting the function of lateral interactions by selectively engaging and/or inhibiting horizontal connections in an adaptive and task-dependent manner^[16]^. Previous work has shown that this spatial integration effect in V1 can be modulated by visual attention demands^[17;18]^.

Active visual behavior, based on saccadic eye movements that bring peripheral targets to the center of vision, requires a highly coordinated system in multiple oculomotor and visual centers of the brain^[19]^. Target selection and movement planning are fundamental for successful saccades. Previous studies have shown that visual attention is directed at the location of future eye fixation in a predictive manner, resulting in presaccadic visual enhancements at the target location^[20]^. These effects are thought to be implemented by top-down feedback from the oculomotor centers to the visual cortex^[21]^, and enhanced pressacadic responses are observed at many different stages of the cortical visual hierarchy, such as V4^[22]^, MT^[23]^, and V1^[24]^.

Multiple behavioral studies have indicated that the spatial spread of presaccadic selection is determined by the visual context^[25–28]^. Therefore, it is plausible that the mechanism underlying such modulation also involves lateral interactions. In this study, we investigated whether lateral interactions are modulated during saccade preparation by testing the effects of lateral flankers on the detection of a peripheral saccade target.

## MATERIAL AND METHODS

### Participants

A sample of 30 participants completed the experiment (19 female, mean age =25.9, age std = 8.3). We excluded data at the participant and condition levels, due to problems in eye-tracking or threshold estimation (see supplemental material for details). All participants were naive to the purposes of the study and had normal or corrected-to-normal vision. This study was approved by the Research Ethics Committee of the Institute of Biosciences - University of São Paulo (Registration Number:25333219.5.0000.5464), and all participants gave their written informed consent before data collection. Participants received a monetary aid of BRL per session to cover personal costs.

### Apparatus

The experimental software was implemented in MATLAB (MathWorks, Natick, USA), using the Psychophysics^[29]^ and EyeLink toolboxes^[30]^. Visual stimuli were presented at a viewing distance of 57 cm on a VIEWPixx monitor (resolution = 1920 × 1080, refresh rate = 120 Hz) (VPixx Technologies, Saint-Bruno, Canada). Eye dominance was determined using a hole-in-card test^[31]^. The dominant eye position was recorded using an EyeLink 1000 infrared eye-tracking system (SR Research, Ottawa, Canada) at a sampling rate of 1000 Hz. Manual responses were recorded through a ResponsePixx 5-button response box (VPixx Technologies). The participants performed the experiment in a dimly lit room with their heads stabilized by a chin rest.

### Experimental procedure

Participants performed a visual detection task (Figure 1A) in two sessions: a saccade session and a fixation session. Eye position was recorded during the entire experiment using an infrared eyetracking system. In both sessions, the task was to detect the presence of a low-contrast target Gabor briefly shown to the right or left of central fixation.

**Figure 1.**
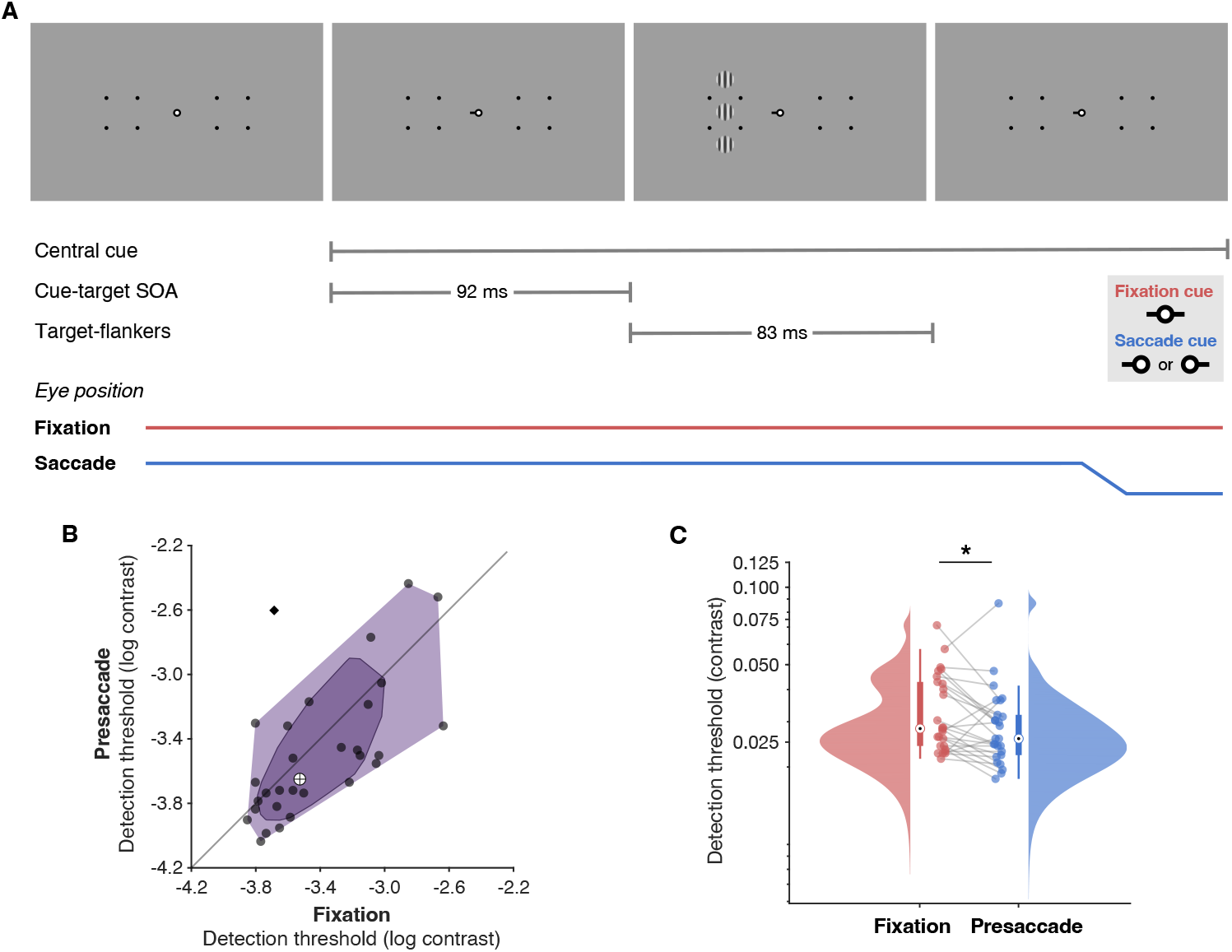
Experimental paradigm. **A**. In each trial, a visual cue appeared at the central fixation point, and after a fixed interval of 92ms (cue-target SOA), the target-flankers stimuli were presented at the right or left hemifield for 83ms. Participants' task was to report whether the target was present (yes/no detection). In fixation trials, the cue was always neutral, pointing towards both hemifields. In presaccade trials, the cue was always predictive of the target location and indicated the direction of the saccade to be executed. **B-C**. Target-only detection thresholds. **B**. Data points represent presaccade and fixation target-only thresholds (log-scaled) from individual participants, and the cross indicates the overall median. The diamond marks an outlier. **C**. Colored clouds represent the density distribution of target-only thresholds. Each boxplot extends to the 25th and 27th percentiles, and the central dot indicates the median. Connected points represent individual participants. The outlier data is not included.

At the start of a trial, participants fixated a central point until a stable fixation within a 2 dva window was detected for one second. Four dot placeholders (diameter:0.3 dva) were presented in a square arrangement (2.25^2^ dva) around the target locations in both hemifields during the entire trial. After a random interval (250-500 ms), a visual cue was presented centrally and remained on the screen until the end of the trial. The cue consisted of a black horizontal line (length: 0.5 dva, width:0.2 dva) pointing right or left (valid cue) or both (neutral cue).

In the saccade session, participants were instructed to make a saccade as fast and as precise as possible to the center of the placeholder at the side indicated by the cue. Visual targets were always presented at the same location indicated by the central cue (i.e. 100% validity). After 92 ms from cue onset, a target Gabor was presented for 83 ms at the right or left hemifield. Participants were instructed to respond whether they had detected the target by pressing one of two response buttons (green button = yes; red button = no). The fixation session had a similar task structure, but the central cue was always neutral, and participants were instructed to maintain central fixation throughout the trial.

The Gabor target could be presented alone or accompanied by two Gabor flankers (Figure 2A). Flankers were positioned in vertical alignment with the target (above and below) at various distances (4,8, or 16 λ). Target and flankers had the same size (diameter: 2 dva) and spatial frequency (3 cycles per degree). The target was always vertically oriented, whereas the flankers’ orientation could be also vertical, in the iso-oriented condition, or rotated by 90° relative to the target, in the orthogonal condition.

**Figure 2.**
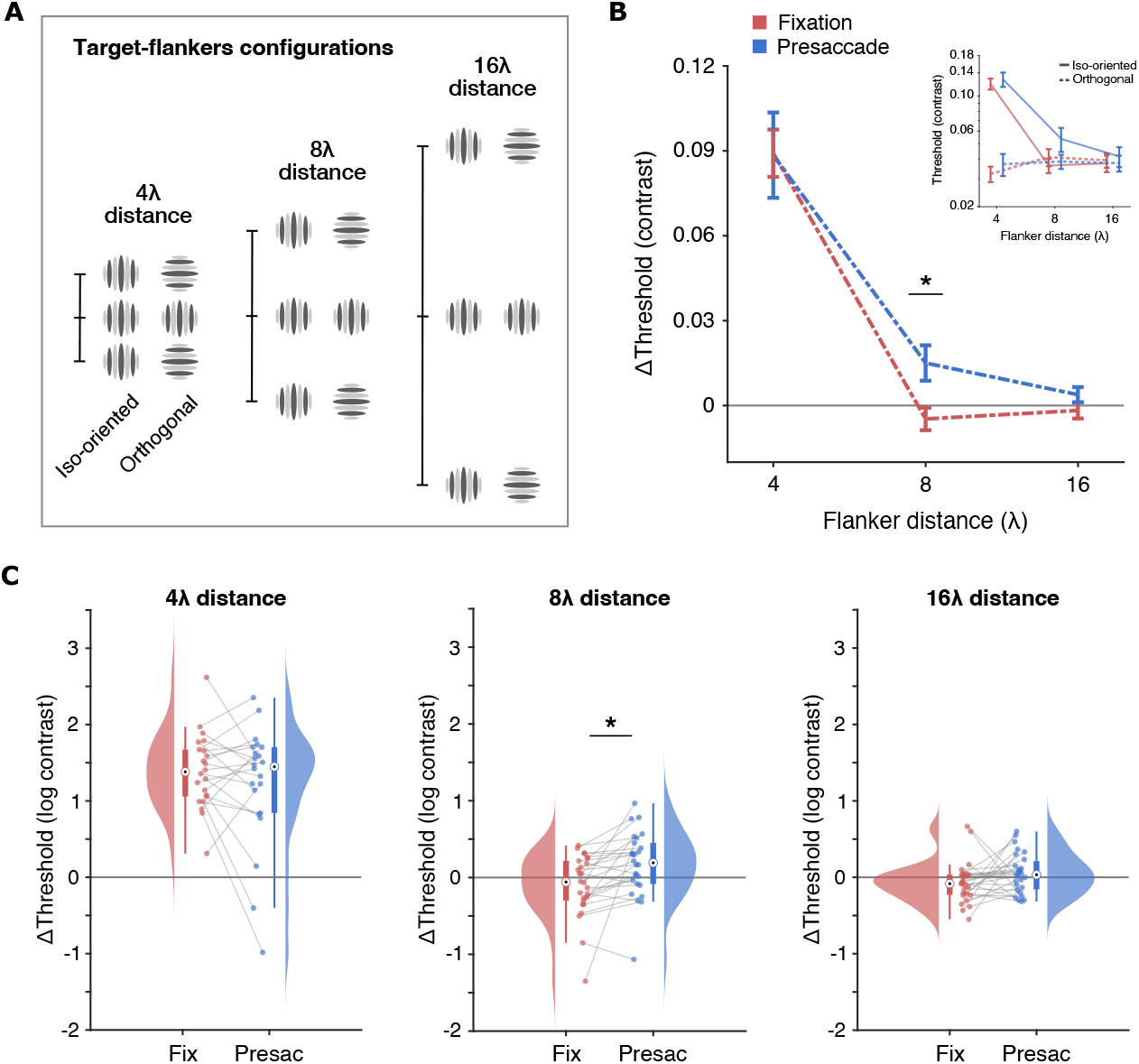
Presaccadic effect on lateral interactions. **A**. Representation of the target-flankers stimuli used in the experimental conditions. **B**. Comparison of thresholds obtained in iso-oriented and orthogonal configurations across the three flanker distances, separately for fixation and presaccade presentations. **C**. Distribution of iso-oriented versus orthogonal (Δ) threshold differences (log-scaled) in fixation and presaccade presentations, separately for each flanker distance. Each boxplot extends to the 25th and 75th percentiles, and the central dot indicates the median. Connected points represent individual participants. Values above zero indicate iso-orientation suppression and values below zero facilitation. **D**. Comparison of fixation and presaccade Δ thresholds across the three flanker distances. Variance bars represent the standard error.

Flankers were presented using a fixed contrast of 0.6 Michelson contrast units (MCU), while the targets’ contrast varied on a trial-by-trial basis according to a staircase procedure, starting at 0.1 MCU. A staircase followed a 3-down/1-up rule, with a step size of 0.1 log MCU, estimating a target contrast level for ≈ 79% detection performance^[32]^. A staircase consisted of up to five 28-trial blocks, and terminated after 140 trials or 12 reversals (whichever happened first).

In each fixation and saccade session, participants performed one staircase per condition: six target-flankers conditions (three distances x two orientations) plus one target-only condition (no flanker). The sequence of conditions was randomized for each participant and session. The sessions lasted approximately 60 minutes, and participants were allowed to take rest breaks between staircases.

### Data analysis

We conducted statistical tests based on linear mixed models fitted to the data. Different models were built for each analysis, as described in detail in the Supplemental Materials. The effects of the factors on the dependent variables were evaluated by performing ANOVAs on the models. Threshold values were given in log-scaled MCU in all statistical analyses.

## RESULTS

### Presaccadic enhancement of target detection sensitivity

First, to investigate whether saccade preparation modulated contrast sensitivity at the target location, we considered the detection thresholds of the target stimulus presented without flankers (target-only condition) (Figure 1B-C). When probed during eye fixation, mean thresholds were 0.034 MCU, and during saccade preparation, mean thresholds were 0.030 MCU. An ANOVA on target-only thresholds and Session (fixation / presaccade) as a factor (Model S1) revealed that detection thresholds were significantly lower for presaccadic targets compared to fixation (F(1,25) = 6.16, p = .020). Thus, we confirmed the classic findings that contrast sensitivity is enhanced for isolated targets presented during saccade preparation.

### Presaccadic effect on lateral interactions

To assess the effects of saccade preparation on lateral interactions, we estimated the perceptual impact of flanking elements on visual targets presented during the presaccadic period or under fixation (Figure 2B-D). Specifically, we used a psychophysical measurement of lateral interactions based on delta thresholds, where the thresholds obtained with iso-oriented flankers were subtracted by the thresholds obtained with orthogonal flankers. Positive delta thresholds indicate iso-orientation suppression, while negative values indicate facilitation.

An ANOVA performed on delta threshold values with Flanker Distance and Session as factors (Model S3) revealed a main effect of flanker Distance (F(2,23) = 49.04, p < .001), no effect of Session (F(1,72) = 1.86, p = .176), but an interaction effect between the two factors (F(2,72) = 5.62, p = .005). Post hoc analyses indicated that only 4 λ conditions were significantly different from zero, with positive delta threshold values in both presaccade (mean =0 088. MCU, std error = 0.015 MCU; t(29) = 9.12, p < .001) and fixation (mean = 0089. MCU, std error =0 008. MCU; t(29) = 10.53, p < .001) sessions, suggesting general iso-orientation suppression at the 4λ distance.

Interestingly, pairwise comparisons on the interaction indicated significant effects of saccade preparation only at the 8λ distance. In particular, 8λ delta thresholds were higher in the presaccade compared to fixation (mean = 0.020 MCU, std error =0.007 MCU; t(72) = 3.02, p = .003), suggesting that saccade preparation increased the suppressive effects of iso-oriented flankers over an 8λ distance from the target. This difference was not observed in distances of λ (mean = -0.0007 MCU, std error =0.014 MCU; t(72) = 1.74, p = .086) and 16λ (mean = 0.006 MCU, std error = 0.005 MCU; t(72) = 1.32, p = .192).

### Presaccadic effect on lateral interactions is not due to saccade statistics

The results of the detection task suggest that saccade preparation modulates the effects of lateral interactions. However, these results could potentially be explained by differences in the spatial and temporal characteristics of saccades that could influence the effectivity of presaccadic modulations between target-flanker conditions. Thus, further analyses were conducted to investigate whether variations in saccade properties could account for the observed effects on delta thresholds.

Across participants and conditions, the median saccade latency was 248 ms. Given that the interval between cue onset and target offset was fixed at 175 ms, the median presaccade interval was 73 ms. The saccade endpoints indicated that, in general, participants accurately directed their eyes to the target region. The vertical amplitude of saccades was near zero, with a median of -0.07 dva, while the horizontal amplitude suggested a slight undershoot from the target center ( 4 dva), with a median of 3.61 dva, but was still within the target region (2^2^ dva) in most cases (Figure 3). Furthermore, participants kept their gaze around the screen center during peripheral target presentation, with eye positions falling near the central point in both horizontal and vertical axes, with a median of -0.003 dva and -0.04 dva respectively.

**Figure 3.**
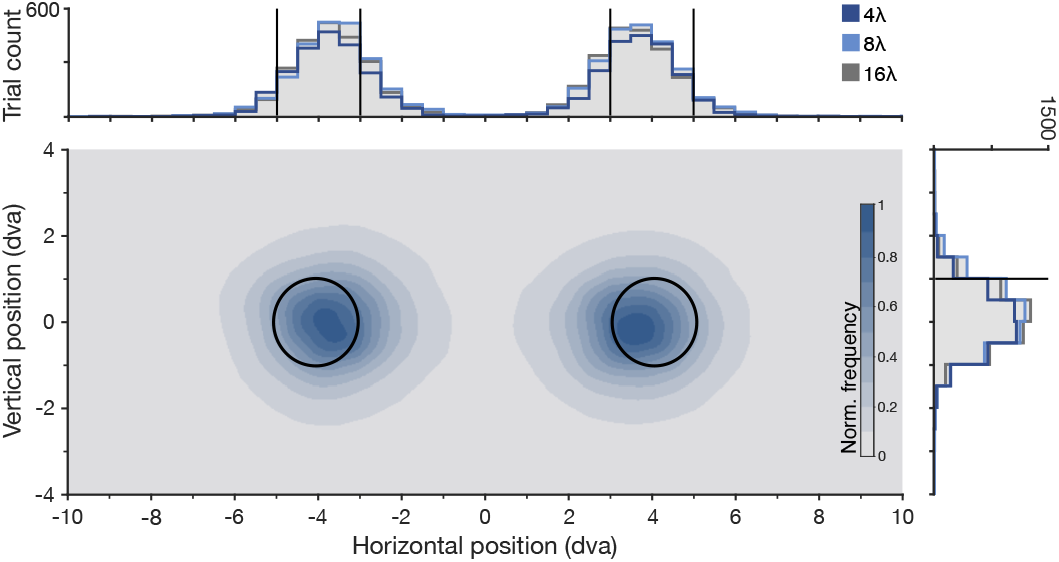
Accuracy of eye movements. Distribution of saccade landing points in the vertical and horizontal axis. Overlapped histograms represent the three flanker distances. Solid black lines in the histograms and black circles in the heatmap indicate the target locations.

An ANOVA conducted on delta thresholds as the dependent variable, with Flanker Distance and median Presaccade Interval as factors (Model S4), showed a significant effect of distance (F(2,51) = 21.57, p < .001), no effect of interval (F(1,45) = 1.87, p = .178), and no interaction between the factors (F(2,57) = 0.64, p = .531). Next, we considered the effect of saccade accuracy, computed as the distance between saccade endpoint and target center. An ANOVA with delta thresholds as the dependent variable, and Flanker Distance and median Horizontal Accuracy as factors (Model S4), revealed a significant effect of distance (F(2,48) = 32.04, p < .001), no effect of horizontal accuracy (F(1,30) = 0.02, p = .886), and no interaction between the factors (F(2,50) = 2.35, p = .105).

A similar ANOVA on delta thresholds, but including the factor of Vertical Saccade accuracy in addition to Flanker Distance (Model S4), showed a significant effect of distance (F(2,47) = 36.34, p < .001), no effect of vertical accuracy (F(1,28) = 0.40, p = .533), and no interaction between the factors (F(2,50) = 0.53, p = .588). Together, these results indicate that the presaccadic effect seen at the flanker distance of 8λ cannot be explained by saccade statistics.

## DISCUSSION

We investigated whether preparing a saccade toward a peripheral visual target modulates the perceptual effects of lateral interactions. Our findings revealed an increase in the suppression of flanked targets presented during the presaccadic period. Specifically, saccade preparation enhanced the suppressive influence of iso-oriented flankers located at 8λ from the target. At 4λ, iso-oriented flankers induced a strong suppression during both presaccadic and fixation conditions. These results suggest that saccade preparation increased the spatial reach of suppressive lateral interactions to at least an intermediate distance of 8λ.

One interpretation of our findings is that saccade preparation modulates spatial integration around the saccade target. In our study, this modulation is observed as an increase in lateral suppression during the presaccadic period. This result is consistent with psychophysical evidence showing an increase in spatial pooling of features surrounding a saccade target^[27;28]^. Although it might seem counterintuitive, an increase in lateral suppression to peripheral targets plays an important role in visual processing. Several studies have supported the idea that in peripheral vision, suppression by iso-oriented surrounds could contribute to masking uniform and redundant regions, allowing more prominent orientation differences to stand out^[9, 11;]^. Favoring the detection of salient regions in the periphery could be more useful in guiding visual behavior and attention, aiding the selection of potential targets for a more detailed inspection by the fovea^[10]^.

It is well known that covert attention can modulate lateral interaction effects^[15]^. Flevaris and Murray^[33]^, for example, have shown facilitation or suppression in V1 BOLD responses, depending on the spatial focus of attention. Directing attention to iso-oriented flankers increased responses in regions representing the center stimulus while directing attention to the central stimulus reduced this response^[33]^. This dissociation was explained by a model combining feature-based attention with response normalization. According to the authors, attention to the flankers increases V1 responses to the center due to the automatic spread of feature-based selection. Conversely, attention to the center would spread more to iso-oriented flankers than to orthogonal ones, thereby increasing the effects of iso-orientation suppression. Interestingly, a recent study has shown that presaccadic perceptual enhancement can spread to non-target items, but only when their features can be grouped with the target’s, following Gestalt principles^[26]^. In our experiment, we observed an increase in suppressive interactions with iso-oriented stimuli during saccade preparation. This could be explained by a presaccadic shift of attention to the target being accompanied by an automatic selection of the iso-oriented flankers, which resulted in a stronger suppression over the central target.

There is a growing body of evidence suggesting that saccade preparation can modulate spatial integration in a time-dependent manner^[34]^. Neupane and colleagues^[35]^ have shown that during earlier stages of saccade planning, neurons in area V4 start responding to input present at their future receptive field location (i.e. forward remapping). However, in later time points, closer to the saccade onset, the neurons’ RFs shift towards the location of the saccade target (i.e. convergent remapping). This increase in the density of neuronal spatial representation could enhance integration around the saccade target. Indeed, Buonocore and colleagues^[27]^ found that the spatial pooling of features surrounding a saccade target was strongest near the saccade onset. Because we used a fixed cue-target interval, contrast sensitivity thresholds were estimated on data pooled from the entire presaccadic period. Future studies could try to dissociate periods of spatial integration and segregation by investigating the time course of the effects of lateral interactions relative to saccade onset.

## ACKNOWLEDGMENTS

This study was supported by the São Paulo Research Foundation (FAPESP), Process Numbers 2017/10429-5, 2018/16635-9, 2019/15213-6, and 2020/09284-5, and by the IDOR/Pioneer Science Initiative (www.pioneerscience.org). We would like to thank Chris Lewis for his helpful comments on the manuscript, the volunteers for taking part in our experiment, and lab colleagues for the discussions throughout the development of this study.

