## Supplemental Materials for "Presaccadic modulation of lateral interactions"

#### Threshold estimation

Contrast thresholds of target detection were determined for each experimental condition using 1-up/3-down staircases[1]. Initial contrast was set to 0.1 Michelson units, increasing by 0.1 log units with each incorrect response (decreasing difficulty) and decreasing by the same amount with three consecutive correct responses (increasing difficulty).

A staircase consisted of up to five 28-trial blocks, and terminated after 140 trials or 12 reversals, whichever happened first. Contrast thresholds, corresponding to 79% of correct responses, were computed by averaging the contrast values in the last 6 reversals.

In each session, participants performed a total of 7 independent staircases for each target-flankers configuration. All the blocks of one condition were performed in sequence. The order of conditions was randomized among participants and sessions. The sessions lasted approximately 60 minutes, and participants were allowed to take breaks between staircases.

The saccade and fixation sessions were held with a minimum of one and a maximum of 14 days apart (mean = 6.9 days), and their order was randomized between participants. The eye tracking was calibrated at the start of each session, and a drift correction check was performed before each block.

Participants were allowed to register their response of target detection anytime after the saccade offset. The position of the response box was switched between participants, so half of them pressed the green ("yes") button using their right hand, and the other half used the left hand. This arrangement was kept the same in the two sessions of each participant.

#### Catch trials

To introduce uncertainty about the presence of the target in the detection task, 15% of total trials in each staircase were catch trials, which did not contain the target stimulus, only the two flankers. Catch trials were ignored in the staircase procedure.

#### Stimuli timing

The timing of stimuli presentation aimed to minimize the possibility that participants directly foveated the target in the saccade session, while also approximating the interval of oculomotor preparation where presaccadic effects can be observed[2]. Specifically, the interval between target offset and saccade onset is when any presaccadic effect can occur. Therefore, the asynchrony between cue onset and target onset was defined based on the expected saccade latency, according to the latency data obtained in a pilot study, aiming to present the target stimulus just before saccade initiation. In addition, participants received feedback on their median saccade latency after each block in the saccade session.

#### Aborted trials

In the saccade session, if no eye signal was detected within a 4 dva window around the target location up to 400 ms after cue onset (saccade latency > 400 ms), the central fixation point turned red and the trial was aborted. This procedure minimized the likelihood of participants directly fixating the target, which would prevent the results from being correctly interpreted as presaccadic effects.

An additional trial was included immediately after an aborted trial, using the same target-flankers configuration and target contrast level as the aborted trial but with 50% probability of the presentation hemifield and 50% probability of being a catch trial.

#### Procedural training

The first experimental session always started with a short training in which participants were familiarized with the task procedure. During training, targets were easily visible, presented at a fixed contrast level of 0.3 Michelson units, or were absent (50% probability). Targets and flankers were presented at different random orientations (0°-180° range, 20° steps), without any orientation (iso/orthogonal) relationship. The three target-to-flankers distances ($4\lambda$, $8\lambda$, $16\lambda$) and the target-only configuration were included.

The training trials were presented in mini blocks (4 trials) separately for each condition. The entire training had a maximum of 48 trials and participants could choose to end it anytime after the first 16 trials (1 mini block per condition). Participants received verbal instructions on how to perform the task and were able to ask questions anytime during training.

#### Data removal

When necessary, data were excluded at the level of participants and conditions. Due to the nature of the staircase procedure, it was not possible to remove individual trials after the experiment had been performed.

Data from two participants were excluded in all conditions due to problems with eye-tracking calibration and/or register, reflecting in tracking errors in more than 50% of the trials (failures in detecting the eye position at the target location). These two participants were also identified as outliers in a boxplot analysis of the frequency of these errors across conditions.

We visually inspected all individual staircases to identify possible problems in the staircase procedure and failures in convergence. If a staircase problem was identified, data were excluded from both conditions (iso-oriented/orthogonal) of the same target-to-flankers distance, since our main analysis was based on the difference between iso-oriented and orthogonal thresholds in each distance.

Data from three participants in the $4\lambda$ conditions were excluded because responses did not lead to the minimum of 6 reversals required for estimating thresholds. In addition, extremely low thresholds were observed in the iso-orientation $4\lambda$ condition of two participants, and in the iso-orientation $8\lambda$ condition of one participant. In fact, these thresholds reached the floor level of contrast in the presentation setup (0.001 Michelson units). These occurrences possibly result from a filling-in effect that leads to a high rate of false alarms when detecting a target between two iso-oriented flankers [3;4].

Finally, data from one participant were excluded in the target-only conditions for being identified as an outlier in the bagplot method [5].

Statistical analysis was carried out without the removals, with data from 22 participants in the $4\lambda$ conditions, 26 in the $8\lambda$ conditions, 27 in the $16\lambda$ conditions, and 26 in the target-only conditions.

### Saccade detection

Saccades were identified in the eye-tracking data using a saccade detection algorithm implemented in MATLAB [6;7]. Parameters included a minimum saccade duration of 8 data samples (8 ms) and a velocity threshold of 5 dva/sec. Velocities were determined using the moving average method. Eye-tracking data were analysed within a time window of 500 ms after cue onset.

### Statistical analyses

Statistical analyses were performed in R Statistical Software using RStudio [8;9] and the packages lme4 [10] and emmeans [11]. Statistical tests were based on linear mixed models (LME), which are adequate for unbalanced datasets and dependent repeated measures, and can also incorporate the effects of inter-subject variability as random variables.

We built LMEs with dependent variables and factors as specified below for each analysis, and participants as a random effect. The effects of the factors were evaluated by performing ANOVAs on the models, with degrees of freedom determined by the Kenward-Roger method. An alpha of 5% was adopted for all significance tests. Thresholds were converted to log-scaled contrast units.

#### Presaccade effect on target-only detection thresholds

For behavioral analysis, we considered as dependent variables: (1) detection thresholds; and (2) delta thresholds. Thresholds were converted to log-scaled contrast units. For computing delta thresholds, the thresholds obtained with iso-oriented flankers were subtracted by the thresholds with orthogonal flankers.

First, we assessed the presaccadic effect on the detection thresholds of an isolated target using a model with the factor of session (presaccade / fixation), and including random intercepts for participants, following the equation:

$$threshold \approx session + (1 \,|\, participant) \qquad \text{(S1)}$$

#### Presaccade effect on delta thresholds

Next, we investigated the effects of lateral interactions more closely using delta thresholds as estimates of these interactions in visual perception, similar to previous studies [12;13]. By using delta thresholds, the effect of iso-oriented flankers is defined in relation to similar target-flankers stimuli that only vary in orientation, rather than to a target-only stimulus.

Delta thresholds were computed separately for each flanker distance and session condition, by subtracting the thresholds obtained with iso-oriented flankers from the thresholds with orthogonal flankers, as:

$$\begin{aligned} \Delta threshold = (threshold\ iso\ oriented) - \\ (threshold\ orthogonal). \end{aligned} \qquad \text{(S2)}$$

The model included delta thresholds as the dependent variable, flanker distance ($4\lambda$ / $8\lambda$ / $16\lambda$), session (presaccade / fixation) and their interaction as factors, and random intercepts for participants with random slopes for flanker distances, resulting in the equation:

$$\begin{aligned} \Delta threshold \approx session + distance + \\ session : distance + (distance \,|\, participant) \end{aligned} \qquad \text{(S3)}$$

Post-hoc contrasts were based on pairwise comparisons among the estimated marginal means for the simple effects of distance and session. Correction of p-values for multiple comparisons was performed using the Tukey method.

**Influence of saccade characteristics on thresholds**

For the eye tracking data analysis, we investigated whether differences in saccade execution between conditions could explain the observed effects on delta thresholds. Of particular interest was the investigation of the presaccadic intervals between target stimulus offset and saccade onset, indicating how long before the saccade the target was presented, and the accuracy of saccade endpoints, indicating how far from the target location the eyes landed.

We constructed models with delta thresholds as the dependent variable, an eye metric (median presaccadic interval or saccade accuracy), flanker distance and their interaction as factors, and random intercepts for participants, according to the equation:

$$\begin{gathered}\Delta threshold \approx eye\,metric + distance + \\ eye\,metric : distance + (1\,|\,participant)\end{gathered} \tag{S4}$$

Post-hoc analyses were performed on fitted slopes of the continuous eye metric variables for each flanker distance. Pairwise comparisons for simple effects of distance were computed based on the estimated slopes, and p-values were adjusted for multiple comparisons with the Tukey method.
